# Complexity-Entropy maps as a tool for the characterization of the clinical electrophysiological evolution of patients under pharmacological treatmentwith psychotropic drugs

**DOI:** 10.1101/164236

**Authors:** J. M. Diaz, D. M. Mateos, C. Boyallian

## Abstract

In the clinical electrophisiologic practice, the reading and comparing electroencephalographic (EEG) recordings some times is insufficient and take to much time. That is why in the last years it has begun to introduce new methods of EEG analysis, that give a better and faster understanding of the EEG dynamics and allow a rapid intervention in the patient’s treatment. Tools coming from the information theory or nonlinear system as an entropy and complexity have been shown to be a very good alternative to address this problem. In this work we introduce a novel method -the permutation Lempel-ziv complexity vs permutation entropy map. This method was applied to EEG of two patients with specific diagnosed pathologies during respective follow up processes of pharmacological changes in order to detect changes that are not evident with the usual inspection method. Our results show that the proposed method are useful for observing an evolutionary retrospective clinical effects of pharmacological interventions in both patients, and from these, to follow the clinical response to the proposed treatment.

## 1 Introduction

The present work arises from the need to have signal analysis tools that allow a more accurate description of the changes observed in the electrophysiologic time series of patients who perform some kind of treatment with psychotropic drugs. From the perspective of Clinical Neurophysiology the electroencephalogram (EEG) is an instrument of observation and monitoring of the electrophysiologic activity of the brain. It has the advantages of being low cost, possible to be replicated and does not imply greater risk for patients. It may also be used in any age range, from preterm infants to older adults [1], in addition to being highly sensitive to temporal changes in the electrical dynamics of the cerebral cortex. This is why it is considered an adequate instrument for the study of the global electrophysiologic activity of the brain. Working with similar objectives we can mention the [2] and as well as the development [3] in the field of electroencephalography in anesthesiology.

With the idea of developing new tools for the EEG analysis, in this work we introduce two methods coming from Information Theory: The *Permutation entropy* (HPE) [4] and *Permutation Lempel Ziv complexity* [5]. Both methods have been widely used in the analysis of EEG signals, obtaining relevant information which is not possible to get by conventional methods, for example power spectrum or wavelets [6, 7]. Particularly in this article we used a parallel analysis of both methods on EEG signals, presenting the results on a Complexity vs. Entropy map. This type analysis representation has been used in chaos versus noise analysis, text authoring analysis, and electrophysiologic evolution of EEG in chickens, among others [8–10].

The clinical objectives proposed for this study were: a) to shorten the latency in the identification of the pharmacological effects on the global neurophysiological activity of the encephalon. b) To facilitate the evolutionary reading in different instances of a pharmacological treatment and the comparison with the respective control groups in order to enrich the epidemiological approach of clinical neurophysiology. c) Shorten the interpretation time of long-term studies with a large number of channels, which usually increase the time of the care task significantly.

The paper is presented as follows: In section 2 we introduce the notions of Permutation Entropy and Permutation Lempel-Ziv Complexity, and then explain the parallel analysis of both methods on the EEG recording; then we show how the control group was conformed and we present a clinical description of two cases with specific pathologies that were considered contrasting with the control group. In Section 3 we present our main results. Here we apply the complexity vs. entropy map to both, control group and two other cases with specific diagnosed pathologies, and we compare them. The discussions about the result are in section 4. Finally, the conclusions and proposals for further developments are presented in section 5.

## 2 Method

### 2.1 Signals Analysis

With the idea to understand and replicate the analysis used in this work, first we introduce the notion of permutation vectors as a quantifier of the continuous signal, then we explain how to calculate the permutation entropy and the permutation Lempel–Ziv complexity. Finally we summarize the steps to apply this method to EEG recordings.

#### Data quantification

When we have to deal with continuous data such as a electroencephalogams (EEG), magnetoencefalograms (MEG) or electrocardiograms (ECG) recordings, an important issue is how to quantify this data. In the bibliography, there are many methods to discretized continuous series as histograms, binarizations, etc. Here we have used the method introduced by Bandt and Pompe called *permutation discretization* [4]. This method is based in the relative values of the neighbours belonging to the series. More precisely, consider a real-valued discrete-time series *{X*_*t*_*}*_*t≥*0_ assumed to be a state of a multivariate trajectory. Take two integers *d ≥* 2 and τ *≥* 1 and define a trajectory in the *d-*dimensional space as

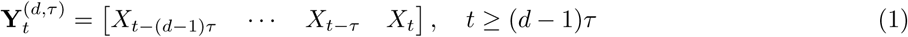

where the dimension *d* is called *embedding dimension* while *τ* is called *delay*.

Now let’s define a vector **Π**^*d,τ*^, called permutation vector as follows. Each component is defined as the relative *t* values of the vector 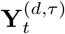 The number of possible motives or patterns (Π^*d,τ*^) is given by the factorial of the embedding dimension *d*! (see Fig.1A). To clarify all this technicalities let’s show how all this works in an example. Suppose we have a continues series such as {*X*_*t*_ = 0.25, 1.5, 3.4, 0.35, 2.2} and take the parameters *d* = 3 and *τ* = 1. The embedding vectors **Y**^(*d,τ*)^ in this case are defined as 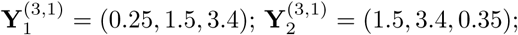, and 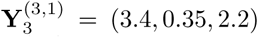 the respective permutation vectors are 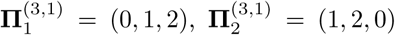 and 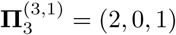 A graphical example of the procedure is shown in Fig.1.

**Figure 1:**
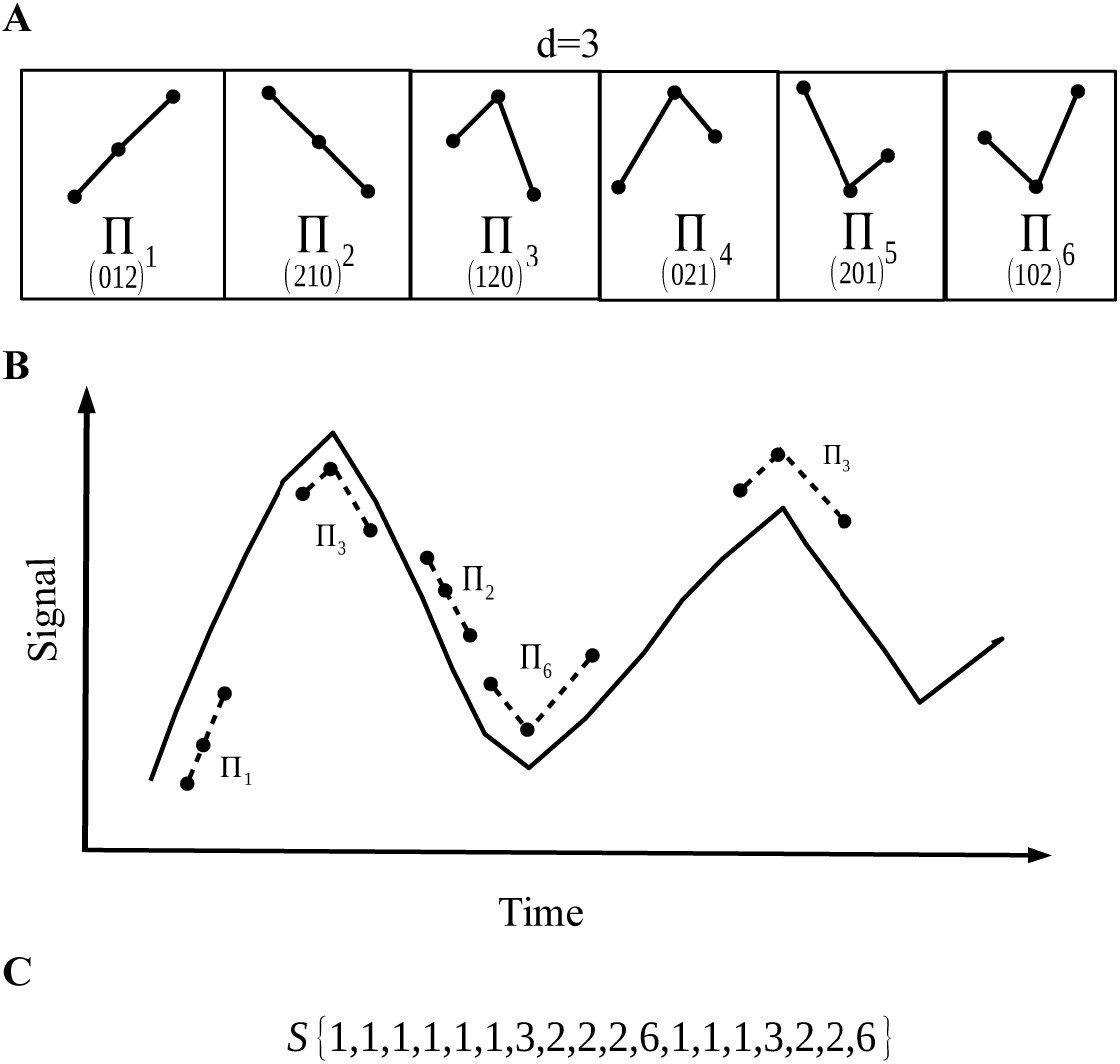
Example of the permutation vector discretization. A) For the parameter *d* = 3 we have d!=6 possible patterns. B) For each data point *t* = 1, *…, n* (*d* 1), is assigned the corresponding pattern, depending on the relative values of the neighbors. C) The quantified sequence to the original signal.

#### Permutation entropy

From a stationary process, the entropy can then be estimated via the frequencies of occurrence of any possible permutation vectors. In their work, Bandt and Pompe defined the *permutation entropy* (PE), as the Shannon entropy of the probability distribution of the permutation vectors, 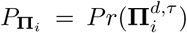 for *i* = 1:::*d*! [4] The normalized permutation entropy is defined as

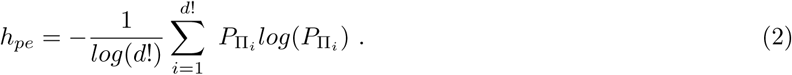

The PE is able to capture the dynamic of the process underlying the data. More precisely, the idea behind PE is that the possible permutation vectors, may not have the same probability of occurrence, and thus, this probability may unveil knowledge about the underlying system.

#### Lempel Ziv Complexity

Now, we will see a different way to analyze a sequence, in this case in not based in the probabilistic of the symbols but in the way that this symbols are repeated along the sequence. It is based in the complexity idea of Kolmogorov- what is the minimal “information” contained in the sequence [11]Lempel and Ziv developed a method to dealing with complexity of a sequence restricting this notion to the “programs” based only on two recursive operations: copy and paste operations. Their definition lies on the two fundamental notions of reproduction and production. Namely, let us consider a finite size sequence *S*_*n*_ = *s*_1_, *…, s*_*n*_ of symbols out of a finite alphabet *A* of *α* letters. Now,

- **Reproduction**: Is a process of reproduction from a sequence *S*_*n*_. It consists in its extension *R*_*n*+*m*_ = *S*_*n*_*Q*_*m*_ where *Q*_*m*_ is a sub-sequence of length *m* of the sequence *S*_*n*_*Q*_*m*_*ϵ* where *ϵ*is the operation of suppression of the last symbol. In other words, there exists an index *p ≤ n* called pointer in the sequence *S*_*n*_ so that *q*_1_ = *s*_*p*_, *q*_2_ = *S*_*p*+1_ if *p < n* and *q*_2_ = *q*_1_ otherwise (the symbol just copied in this case), ect. As an example,*S*_3_ = 101 reproduces the sequence *R*_6_ = 101010 since *Q*_3_ = 010 is a subsequence if the *S*_3_*Q*_3_*ϵE* = 10101, *q*_1_ = *s*_2_ = *r*_2_, *q*_2_ = *s*_3_ = *r*_3_ and *q*_3_ = *q*_1_ = *r*_4_ previously copied. In other words, *SQ* can be reproduced from *S* by recursive copy and paste operations. In a sense, all the information of the extended sequence is in *S*_*n*_.
- **Production**: Is a production operation denoted *S*_*n*_ *→ R*_*n*+*m*_ = *S*_*n*_*Q*_*m*_ consists in reproducing the subsequence *R*_*m*+*n*_*ϵ* from *S*_*n*_. As an example *S*_3_ = 101 produce *S*_3_*Q*_3_ = 101011 but it does not reproduce this sequence *q*_1_ = *s*_2_ = *r*_2_, *q*_2_ = *s*_2_ = *r*_3_ but the last symbol does not follow the recursion, *q*_3_≠*q*_4_. The difference between reproduction and production is that the last letter can come either from a supplementary copy-paste or it can be new.

Now, any sequence can be viewed as constructed through a sequence of production processes, which is called history H. Indeed Ø → *s*_1_ → *s*_1_*s*_2_ →*s*_1_*s*_2_*s*_3_→*…* In this example, starting from the empty string, we have *n* production operations: the size of the history is *n*. However, a sequence has not an unique history. Indeed, in the case where *s*_2_ = *s*_1_, one can reduce the number of production processes since Ø *→ s*_1_ *→ s*_1_*s*_2_*s*_3_ *→…*Then, for a given history H _*i*_ of the sequence, let us define by 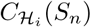 the number of production porecesses in given history. Clearly min {2, *n*}≤*C* H *i* (*Sn*) ≤ *n*. Lempel and Ziv define the complexity of the sequence as the minimal number of production processes needed to generate it:

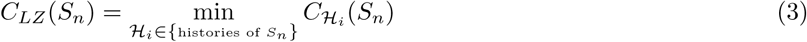

A drawback of this complexity is that it is defined only for sequences of symbols taken in a discrete finite size alphabet. Dealing with “real life” sequences a quantization has to be performed before its use, as done in many of the studies dealing with data analysis via the Lempel–Ziv complexity [12–14]. Using the same idea that in permutation entropy, the alphabet can be taken as the set of permutation vectors 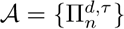 and the alphabet length α = *d*!. This is called *permutation Lempel–Ziv complexity* (PLZC) [5].

Surprisingly, although we are analyzing a sequence from a completely deterministic point of view, it appears that *C*_*LZ*_(*S*_*n*_) sometimes also contains a notion of information in a statistical sense. This make it emerge a relation between the Lempel–Ziv complexity and the Shannon entropy [15–17]. Using this relation, we define the **permutation Lempel–ziv complexity vs permutation entropy map**. The idea behind this approach, is that the analysis of a signal by means of this map, gives us more information than the separate use of each measure. Next we will explain the steps to analyze a signal through the entropy complexity map.

#### Complexity vs entropy map

Now, we will explain the step to analyze the EEG recording data and the representation in the result in the complexity vs entropy map.

- Step 1: Recording and preprocessing the signal. Using band pass filter and notch filter (Fig.2A).
- Step 2: Discretize the raw signal using permutation vector approach (Fig.2B).
- Step 3: Calculate the Lempel-Ziv complexity for the sequence taken from step 2 (Fig.2C).
- Step 4: Take the sequence from step 2 and using a histogram, estimate the probability distribution (PD) of permutation vectors. Then, calculates the Shannon entropy related with this PD.
- Step 5: Finally with the two measures we have the coordinates (*h*_*P* *E*_, *C*_*LZ*_) of the map corresponding to the recorded signal.

**Figure 2:**
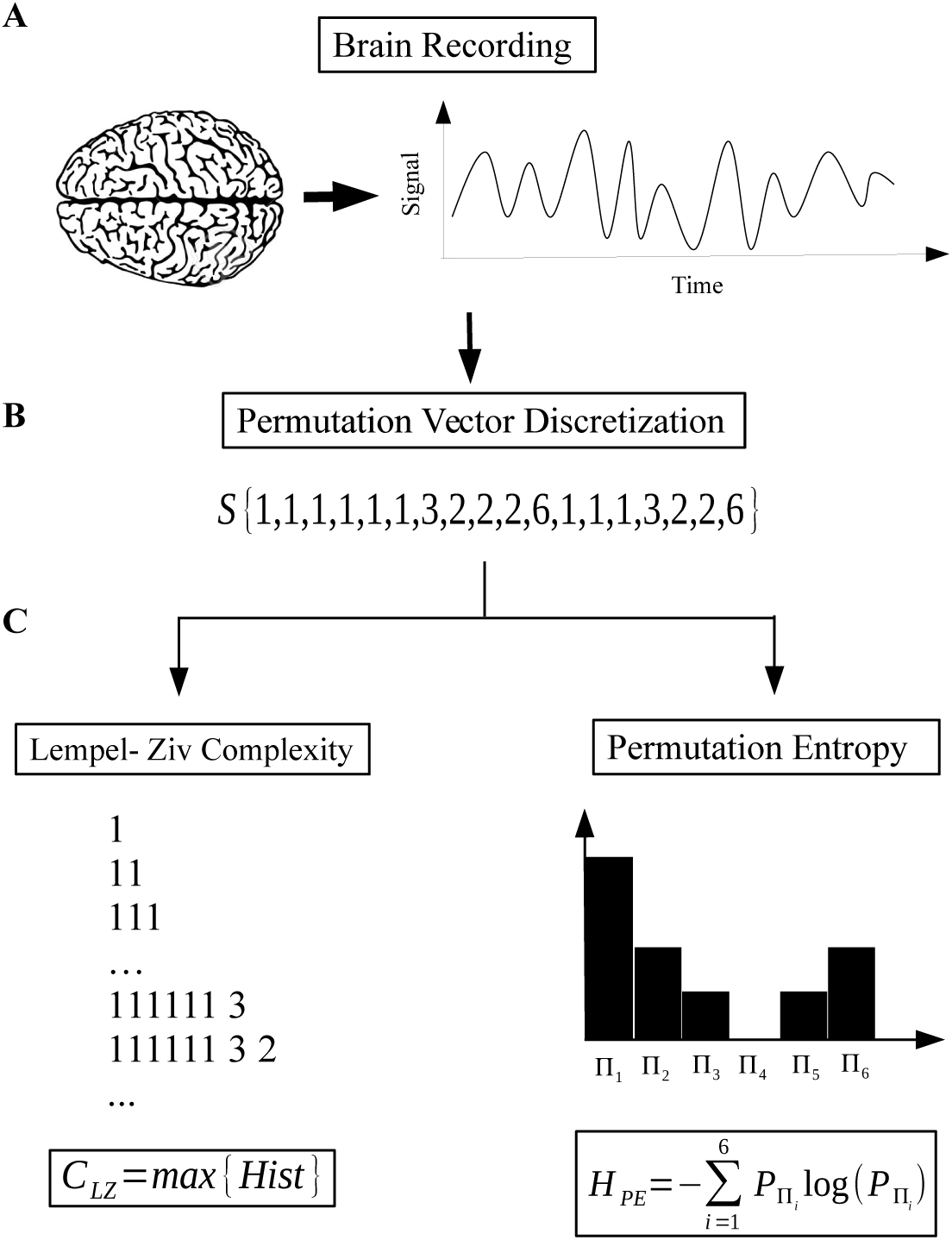
The steps to analyze a recording signal through the complexity vs entropy map. A) Signal recording and preprocessing. B) Quantify the signal using the permutation vector approach. C) Calculate the permutation Lempel ziv complexity and the permutation entropy for the quantify sequence.

To make a statistical analysis, for example determinate the mean value and standard deviation of different channels or for a group of people, it is required to calculate the covariance matrix Σ_*h,c*_. For a group of measure *h*_*P* *E*_{*h*_1_, *…, h*_*n*_} and *C*_*LZ*_{*c*_1_, *…, c*_*n*_}., the matrix entries are given by:

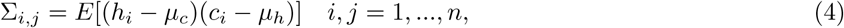

where *E* is the expectation value and *µ*_*c*_, *µ*_*h*_ are the mean value of the entropy and the complexity respectively.

### 2.2 Patients recording

The present work has a comparative, observational and retrospective design between a control group versus two clinical cases of patients with diagnosed pathologies that were studied and followed by more than one year of evolution in a laboratory of clinical neurophysiology. During the study period the patients with diagnosed pathologies followed the therapeutic indications of their treating physician and electroencephalographic studies were carried out in different instances of their treatment. In both cases these patients were under treatment with psychotropic drugs. The electroencephalographic studies were analyzed retrospectively and compared to the parameters obtained after applying the same methods to the control group.

#### Technical procedures and clinical description

For the conformation of the Control Group, *N* = 20 EEG recordings were selected from a database of clinical neurophysiology laboratory (IACCo). Inclusion-exclusion criteria were: a) records for adult individuals (18-50 years) with no medical history. b) Should not be taking psychotropic medication at the time of the study. c) The study should have been performed without facilitator medication like chloral hydrate and / or similar. (d) EEG records had to be reported as normal pathways by the specialist physician, (e) records should not contain non-physiological artifacts. (f) Studies that conformed the control group should have a similar duration. (h) The studies of the control group should be performed under equal conditions (closed eyes and similar number of samples and electrodes) than the pathological cases studied.

Each record analyzed here was performed according to the standards of the International Federation of Clinical Neurophysiology [18], following the international system 10-20 for the placement of electrodes. The records are presented in monopolar montages, with bimastoid reference for a total of 20 channels. The signal was taken with a low resolution sampling of 65*Hz*, with conventional hospital use equipment, according to local regulations. The number of the individuals was *N* = 20 and the mean ages was 33*±*10 of both sexes and the duration of the records was 37*±*9 minutes for all cases in the control group. All the information about the Control group resume in table 1

**Table 1:** Control Group Information

| Subject | Time Record (min) | Segment time analyzed (min) | Gender | Age |
| --- | --- | --- | --- | --- |
| 1 | 60 | 44 | M | 24 |
| 2 | 60 | 49 | M | 20 |
| 3 | 60 | 44 | M | 31 |
| 4 | 60 | 44 | M | 48 |
| 5 | 60 | 44 | M | 25 |
| 6 | 60 | 48 | M | 38 |
| 7 | 30 | 30 | F | 27 |
| 8 | 30 | 30 | F | 28 |
| 9 | 30 | 30 | M | 19 |
| 10 | 60 | 30 | F | 32 |
| 11 | 30 | 30 | F | 44 |
| 12 | 30 | 25 | F | 40 |
| 13 | 30 | 25 | F | 38 |
| 14 | 60 | 48 | M | 48 |
| 15 | 60 | 39 | M | 34 |
| 16 | 60 | 50 | M | 50 |
| 17 | 60 | 40 | M | 23 |
| 18 | 30 | 30 | M | 23 |
| 19 | 60 | 41 | M | 31 |
| 20 | 60 | 45 | F | 27 |

### 2.3 Clinical description of the Cases

#### Case I

The records in this case correspond to a female patient of 22 years of age, whose clinical diagnosis was Idiopathic Generalized Epilepsy (ICD-10 G 40.3). She had clinical follow-up for a period of two years. During this time, 4 EEG recoding were performed. The references of these studies will be *T*_0_, *T*_1_, *T*_2_ and *T*_3_, respecting the chronological order in which they were realized.

The initial condition from which the case description starts is *T*_0_. This study was performed when the patient was taking 1200*mg/day* of carbamazepine, without any improvement. This situation led to an average of 10 seizures per week. After her first EEG recording (*T*_0_), the pharmacological scheme was changed to carbamazepine 400 *mg/day* and valproic acid 1000 *mg/day*. This change improved the clinical state and the number of seizures reduced to two per month. This improvement persists throughout the first year of follow-up and it is reflected in the following EEG recording *T*_1_ taken 10 month after *T*_0_. Inmmediately after *T*_1_, a new pharmacological scheme was implemented, including Lamotrigine 400 *mg/day* and 1500 *mg/day* of Levetiracetam. This change brought the seizure frequency to a seizure every three months. After 7 months of this new pharmacological scheme, *T*_2_ was recorded. The latter improvement observed in the clinic could not be discriminated by the method of visual inspection of the signal and the neurophysiological report of *T*_2_ was described as an evolution without significant changes. At this point, the dose of one of the drugs of the last implemented scheme was increased. This is, Levetiracetam 2000 *mg/day* and maintaining Lamotrigine 400 *mg/day*. After this change the patient was for more than 6 months without seizures. Seven months later *T*_3_ was recorded. The time elapsed between *T*_0_ and *T*_3_ is approximately of two years of treatment. In all studies there were no seizures during the recording, so the tracings correspond to intercritical records.

Electroencephalographic records 3 were reported as: *T*_0_, Severe Bilateral Disorganization; *T*_1_ and *T*_2_ records were reported as Moderate Bilateral Disorganization, in which no significant differences were found between both registries and a favorable evolution between both in comparison with *T*_0_. The last one, *T*_3_ was described as a Bilateral Disorganization leading to frontal predominance, with a significant improvement in relation to the previous tracings.

#### Case 2

This second case corresponds to a 41-year-old man diagnosed with Narcolepsy (CIE-10 G47.4) and morbid obesity as co-morbidity. In this case three follow-up studies were made, namely *T*_0_, *T*_1_ and *T*_2_ in an overall period of 16 months. Again, the subindices represent the chronological order in which the studies were performed.

At the initial condition *T*_0_, the patient was medicated with Topiramate 250 *mg/day* because of his eating disorder. However this medication provoked no changes in symptoms related with Narcolepsy. After *T*_0_, the pharmacological scheme is changed and modafinil 200 *mg/day* is added. After 9 months with this new scheme (topiramate 250 *mg/day*, plus modafinil 200 *mg/day*), a clinically significant change was achieved in relation to nocturnal rest and a marked decrease in daytime sleepiness, which also improved the patient’s performance in daily life activities. At his point *T*_1_ was recorded and no pharmacological changes were introduced. Seven months later, *T*_2_ was performed just to check the evolution of the patitient. In this last evaluation, the recording time was doubled in order to record sleep activity in the EEG, however the patient kept awake with his eyes closed during the two hours of the study. As expected from its favorable evolution, daytime sleepiness continued to decline as treatment time advanced.

In the electroencephalographic records Fig. 4, *T*_0_ was reported as a Severe Sleep Cycle–Vigil Disorder with a Normal Vigil plot. Here fig.4A, *K* complexes and sleep spindles can be observed, which are characteristic of Phase II, non-REM sleep. These patterns are best presented at the electrodes near the electrode *C*_*z*_. In *T*_1_ (fig.4B) and *T*_2_ (fig.4C) tracings were reported as tracings within normal limits. In the retrospective evaluation between the *T*_1_ and *T*_2_, no significant clinical differences were found. Both traces correspond to closed eyes awake records.

**Figure 4:**
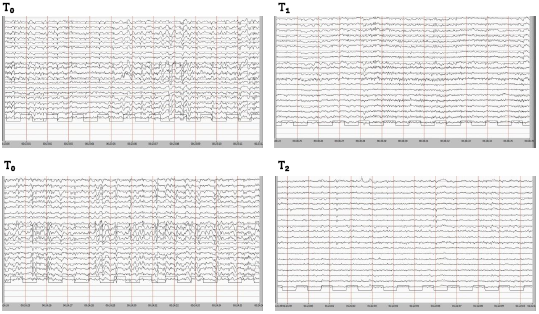
EEG traces belonging to patient diagnosed with Narcolepsy (Case 2). *T*_0_ correspond to sleep paths, in both can see the slow waves between 3 and 5*Hz*. In addition, *K* and spindles complexes, characteristic of phase 2 of the non-REM sleep, are observed. The *T*_1_ and *T*_2_ traces correspond to normal record in waking eyes closed condition. *T*_2_ trace corresponds to the second hour of the monitoring. A trace of low voltage with alpha and theta rhythms is observed in both records.

## 3 Result

In both cases and each of the 20 patients conforming the control group, we analyze the EEG’s under the Permutation Entropy (PE) and the Permutation Lempel–Ziv complexity (PLZC), using the same permutation vector parameters. Every EEG channel were analyzed separately and using the whole recording. The results were shown in an complexity-entropy map. The parameter used for the permutation quantification where *d* = 4, 5, 6 and *τ* = 1, giving similar results. The center of the ellipse corresponds to the mean value of the coresponding studies of the control group and the ellipse itself, to the joint error of PE and PLZC of this group.

For our first patient (case 1), Fig. 5 shows the results of the 4 EEG monitorings performed on the patient in an evolutionary follow-up of a 22-year-old woman diagnosed with Generalized Idiopathic Epilepsy. Here a clear displacement in the plane between the *T*_0_ and *T*_1_ is observed. The interval between the two studies was 10 months and it corresponds to a large decrease in the frequency of seizures.

**Figure 5:**
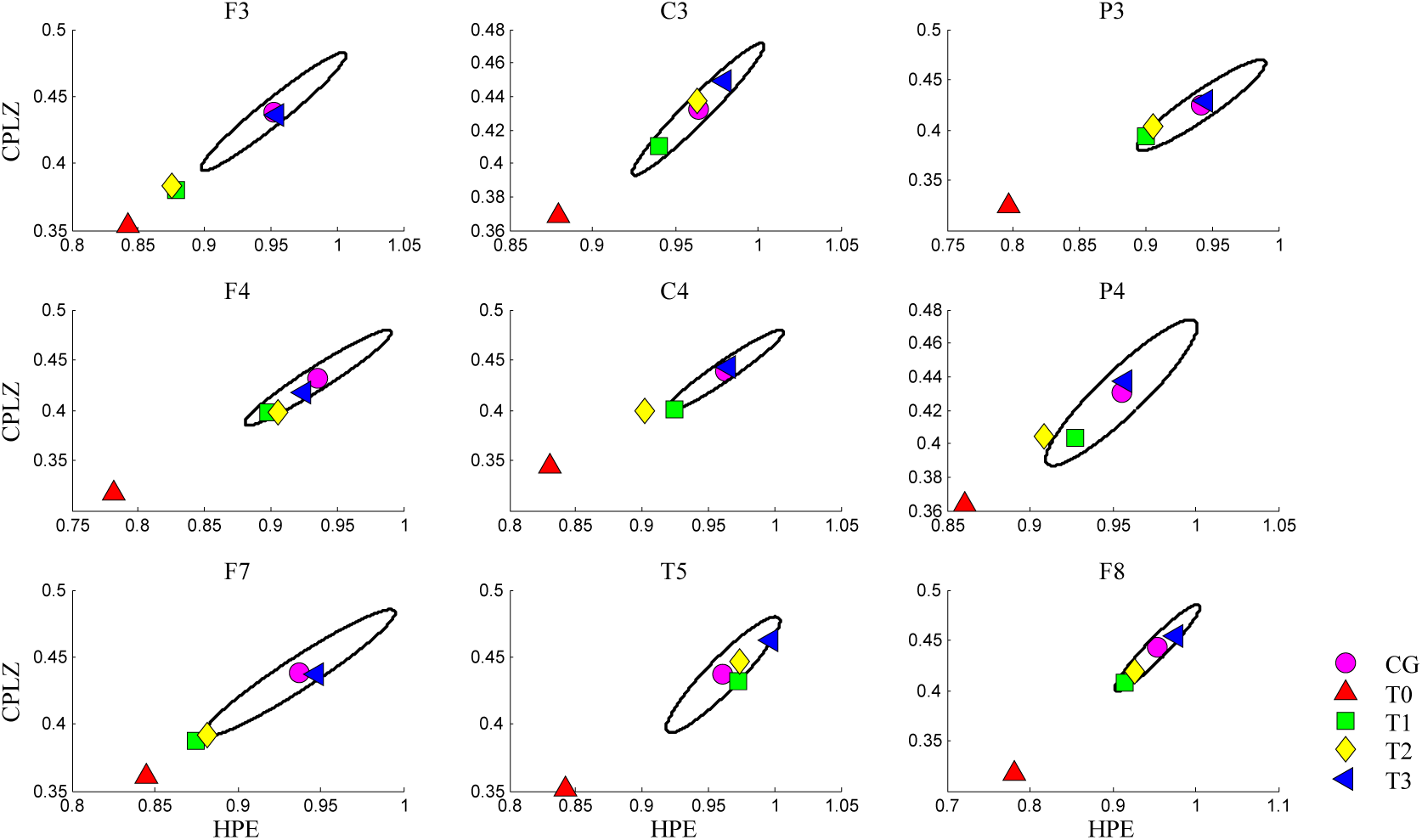
Analysis throw the PE vs PLZC map belonging to the 22 year old woman diagnosed with generalized idiopathic epilepsy. Each subplot shows the values of complexity and entropy in the different 4 stages of her treatment compared to the mean value of a control group and their respective error (the parameters used were *d* = 4 *τ* = 1))

The time elapsed between *T*_1_ and *T*_2_ was 7 months. This corresponds to the change in the pharmacological scheme that sought to prevent the patient from having seizures, not only to reduce their frequency. Here the neurophysiological report of the electroencephalogram by visual inspection found no significant change between them. However, there was an improvement in the clinical aspect of the condition. Finally in *T*_3_ can be observed that the optimization of the pharmacological scheme used until *T*_2_, achieves a new significant change both in the clinical aspect, as well as in the electroencephalographic tracing, which is approaching in many channels to the center of the ellipse that corresponds to the to the mean value of the studies of the control group. The total time elapsed between *T*_0_ and *T*_3_ is two years of treatment evolution. In addition, it is important to note that the 4 pathological studies presented in this case correspond to intercritical studies, since no seizures occurred during the recording.

Figure 3 (case 2) shows the results of the evolutionary follow-up of a 41-year-old male with a diagnosis of Narcolepsy and Morbid Obesity. Here the 3 electroencephalographic monitoring can be observed, comparing the differences between the abnormal study *T*_0_ (Severe Sleep Cycle Vigil Disorder) with *T*_1_ and *T*_2_, which are reported as normal. During the period of time elapsed between both normal records, approximately 90 days, the scheme topiramate 250 *mg/day*, and modafinil 200 *mg/day* was kept. For this reason it is considered that both studies were performed under conditions of pharmacological stability. However, between *T*_1_ and *T*_2_ there is little difference: *T*_1_ lasted one hour and *T*_2_ lasted two hours with a previous sleep deprivation. Calculations for *T*_2_ were analyzed for each hour separately and similar results were found between the two halves, so that the graph shows the values of the complete study (2hs). It is also possible to identify that the values of the plane for the study *T*_0_ are lower in the electrodes near *C*_*z*_. Finally, it should be mentioned that it was from the implementation of the mentioned pharmacological scheme with which the clinical improvement was achieved. This clinical improvement was reflected as a permanent feature of the patient’s functioning in his daily life. This corresponds to the pharmacological stability and the proximity between the values of studies *T*_1_ and *T*_2_, and also to similar results found in both hours of *T*_2_, which showed a normal tracing of wakefulness of closed eyes.

**Figure 3:**
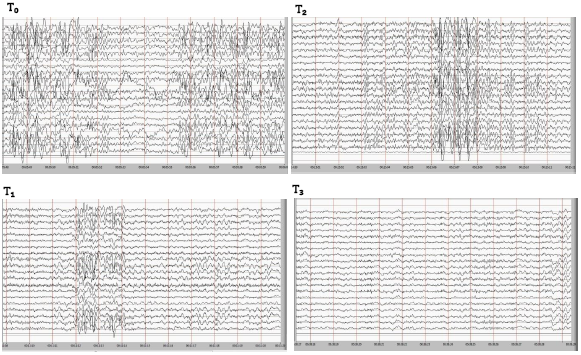
EEG recording example for the four states of treatment (Case 1). *T*_0_ corresponds to the first monitoring performed, in it can be observed that before the time 05: 52 and after the time 5: 55, the discharges occur in both hemispheres and had a duration greater than 10 seconds. In the studies *T*_1_ and *T*_2_, bilateral short-term discharges 2-3 seconds are observed. There is little clinical differentiation between studies *T*_1_ and *T*_2_. In the study *T*_3_ a trace with a normal voltage with a predominance of bilateral slow rhythms in frequency domain is observed. All studies correspond to intercritical records.

## 4 Discussion

The results allow us to compare the values obtained for each channel of an EEG, with the mean value of the values of the control group and the corresponding error (ellipse). This deepens the signal reading, supplementing the traditional method of visual inspection of EEG, which is usually performed on a time scale of few seconds and whose reading is done sequentially.

In the first case the map is used as a tool that allows to show the effectivity of pharmacological treatment in a patients treated with psychotropic drugs. In a pathology such as epilepsy, the EEG signal tend to be more regular. This regularity gives less number of different permutation vectors and this provokes a decrease in the entropy and complexity values. Similar result where shown comparing awake people with people in altered state of consciousness such as sleep, seizure and coma, [19]. Analogous results are shown in other epilepsy syndromes such as absence crises, [20]. In our first case, although there is an interelectrode difference that can be observed in the plane, there is evidence of a common trend in all channels, which shows an approach to the control group, which reflects a change in the functional state of the nervous system that correlates with the favorable evolution after each pharmacological intervention. Moreover, when the visual inspection method of EEG did not shows significant changes between the *T*_1_ and *T*_2_ studies, the pharmacological scheme was changed and it corresponded to an improvement in the clinic. The complexity-entropy plane was able to detect such a change from the global evaluation of the entire record for each electrode. As can be seen in figure 5, 6 of 9 channels shown there show smaller distances to the center of the ellipse in *T*_2_ compared with *T*_1_. This subtlety, that had clinical value, could not be detected by the human eye. Also, by optimizing the dose, a major change is achieved and it visible in *T* 3. EEG records in epilepsy are now described as critical and intercritical records. However, most of the studies that are done in clinical practice are intercritical records, because there is no way to predict a seizure and only in some cases can the crises be reproduced, such as those that respond to sensitization methods, such as intermittent light stimulation, and usually have a genetic or syndromic association that is still unknown [21]. Therefore, a quantitative evaluation of the intercritic records opens the possibility of comparing studies in which it is not possible to perform a semiologic description of the seizure, but from which it is possible to obtain valuable information about the clinical evolution of the epileptic patient.

**Figure 6:**
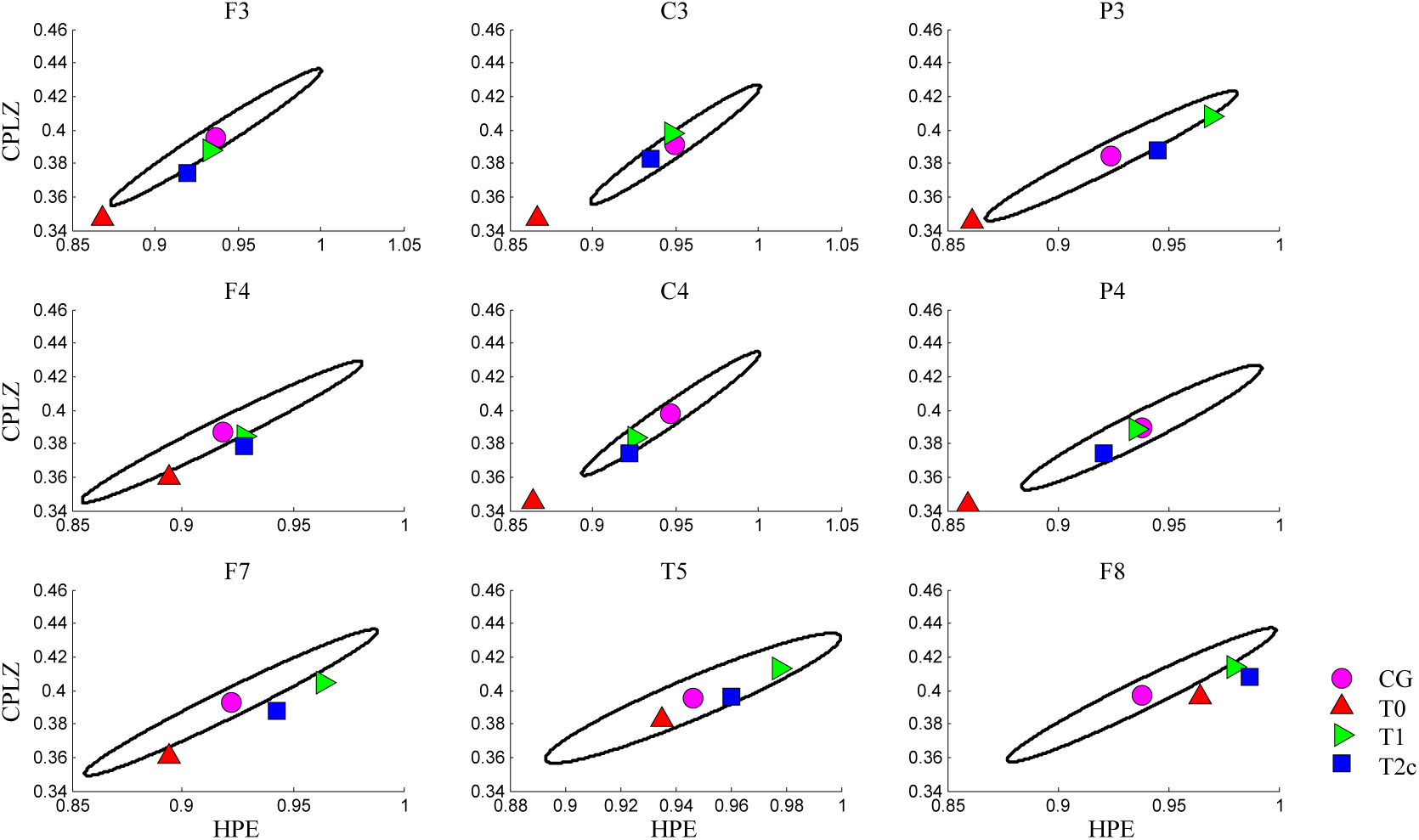
Analysis by complexity vs entropy map of a patient who suffers from Narcolepsy. Each subplot shows the complexity and entropy values in the different 3 stages of his treatment compared to the average value of the control group and its respective error (the parameters used were *d* = 4, *τ* = 2).

In the second case, which corresponds to the 41-year-old patient with narcolepsy, the CLZvsHPE method of analysis is also used. Recall that the EEG sleep pattern is different from the awake pattern. These differences in paths that are observed by the visual inspection method (Fig. 4) are also identified by the CLZvsHPE plane. In the images of *T*_0_ in Fig. 4, some patterns of the characteristic pattern of non-REM sleep phase 2 such as *K*-complexes and splindes ([23, 24]) can be found. During this study, the patient was able to maintain the waking state during very short periods of time and most of the EEG tracing corresponds to a sleep pattern. Here CLZvsHPE values are very low values for most electrodes; however, in daily clinical practice, low values in the plane are unusual for short studies, since most patients usually maintain vigil or enter a light sleep state in records of an hour or less duration. This is related to the low values found in individuals with altered states of consciousness such as those presented in [19].

After the administration of modafinil, the tracings have characteristics normal patterns of the vigil state with closed eyes. In Fig. 4, segments of these studies are shown and correspond to *T*_1_ and *T*_2_. The values found in the CLZvsHPE plane for such studies are similar to the values found in control individuals.

The similarity of these values in both studies correspond to the fact that the patient treated with modafinil managed to stay awake during the whole records. In the channels *F*3 – *C*3 – *F*4 – *C*4 – *F*8 of Fig. 3 it is possible to observe the small difference between the values obtained in the plane for the *T*_1_ and *T*_2_ studies. Note that the time elapsed between this studies is 7 months and *T*_2_ is twice as long as *T*_1_. The approximation of the values of both studies to the center of the ellipse shows an improvement of the patient with respect to *T*_0_. The fact that there is a proximity in the values of the channels between the two studies (*T*_1_ and *T*_2_), indicates certain stability in the improvement, which correlates with the clinical evolution that the picture had.

Note that in this work low values in the plane correspond to pathological clinical conditions or changes in the state of consciousness. [19, 20]. For example, as shown in the first case, the low values correspond to changes found in the signal given by patterns (graphs) of pathological discharges in intercritic studies of a patient with epilepsy. However, in the second case, pathological patterns (graphemes) are not found in the EEG, but physiological patterns of normal sleep that should not appear in a closed-eyes waking study if the patient could maintain alertness during short duration EEG and without facilitating medication.

In this way the CLZvsHPE map would provide new information to methods like MSLT (The Multiple Sleep Latency Test) and polysomnography for the evolutionary and / or diagnostic studies of narcolepsy. In addition, as shown in case one, in the evolutionary study of patients with neuropsychiatric pathologies who are treated with psychotropic drugs relevant information is provided by the plane. The importance of this is found in the fact that there is an individual variation in the response to psychotropic, an aspect that if taken into account would achieve in the future the development of a more personalized drug therapy.

## 5 Conclusion

In this paper show us one application of “PLZC vs PE” map for electroencephalographic interpretation in clinical neurophysiology. This method allows us to look at the changes that occur in long term treatments from a single perspective. Something that the method of visual inspection can not provide us so far, is to be able to compare more than one study in a single image, to obtain a single value for each electrode of the complete signal and to allow the comparison between different electrodes in a same patient.

We applied the analysis to a control group in order to have a statistical reference. This provides an epidemiological approach to the analysis of each clinical case and offers a better understanding for the conventional clinical interpretation of the data. Here we find that for the healthy people the entropy and complexity of signal tend to be maximun and it is a dynamical interpretation of our results. The instrument presented here has been used as a global assessment tool, sacrificing short-term signal changes, that is, its non-stationary behavior. However it is also possible to apply the instrument with sliding windows and retrieve this information if it is considered relevant [19]. As an additional feature, the planes corresponding to each of the 20 electroctodes can be presented following the international 10-20 electrode placement system, which will become familiar with the usual reading for health workers, even if the number of electrodes is greater, namely, 32, 64 or 128. In clinical medicine, longitudinal follow-up of cases is always a daily practice. However, gathering information from several studies (EEG) of the same patient and making an evolutionary reading, is an expensive and complex work.

Finally, and taking up the clinical objectives of this work, we consider that the instrument is useful for the evolutionary follow-up of patients undergoing pharmacological treatment with psychotropic drugs. The epidemiological and statistical approach in electroencephalography is accessible if the control groups can be adjusted by age groups, the latter being especially important in neonatal electroencephalography. For prolonged studies of hours or days of patients hospitalized in electroencephalographic monitoring units, the instrument offers the possibility of reading each recorded signal hourly, which helps to reduce the reading time of such monitoring in the medical care task

In relation to pharmacodynamic studies by means of EEG signals, the spectrogram is one of the most widely used methods in the field of anesthesiology. The instrument presented here was not used to this purpose, however we think that it can add information to the data obtained by the spectrogram and contribute to the investigation in this field. This is part of the work in progress.

